# Statistical significance and biological relevance: The case of *Nosema ceranae* and honey bee colony losses in winter

**DOI:** 10.1101/2022.05.20.492825

**Authors:** Vivian Schüler, Yuk-Chien Liu, Sebastian Gisder, Lennart Horchler, Detlef Groth, Elke Genersch

## Abstract

Managed and wild insect pollinators play a key role in ensuring that mankind is adequately supplied with food. Among the pollinating insects, the managed Western honey bee providing about 90% of commercial pollination is of special importance. Hence, diseases as well as disease causing pathogens and parasites that threaten honey bees, have become the focus of many research studies. The ectoparasitic mite *Varroa destructor* together with deformed wing virus (DWV) vectored by the mite have been identified as the main contributors to colony losses, while the role of the microsporidium *Nosema ceranae* in colony losses is still controversially discussed. In an attempt to solve this controversy, we statistically analyzed a unique data set on honey bee colony health comprising data on mite infestation levels, *Nosema* spp. infections and winter losses continuously collected over 15 years. We used various statistical methods to investigate the relationship between colony mortality and the two pathogens, *V. destructor* and *N. ceranae*. Our multivariate statistical analysis confirmed that *V. destructor* is the major cause of colony winter losses. When using cumulative data sets, we also found a significant relationship between *N. ceranae* infections and colony losses. However, determining the effect size revealed that this statistical significance was of low biological relevance, because the deleterious effects of *N. ceranae* infection are normally masked by the more severe effects of *V. destructor* on colony health and therefore only detectable in the few colonies that are not infested with mites or are infested at low levels.

## Introduction

The basis of human nutrition includes agriculturally grown crops and fruits, many of which are dependent on insect pollination for fruit set, seed production, and yield. Managed and wild insect pollinators therefore play a key role in ensuring that mankind is adequately supplied with food (Aizen et al., 2008; Aizen et al., 2009; Garibaldi et al., 2013). As a result, the health and survival of pollinating insects have attracted increasing public and scientific interest and consequently diseases as well as disease causing pathogens and parasites that threaten pollinating insects have become the focus of many research studies. In terms of pollinating insects, the main focus is on the Western honey bee *Apis mellifera*, which is managed by beekeepers for honey production all over the world and provides 90% of the commercial pollination worldwide (Aizen et al., 2008). In terms of pathogens, the focus is on those that threaten the survival of the managed honey bee colonies. The ectoparasitic mite *Varroa destructor* together with deformed wing virus (DWV) vectored by the mite have been identified as the main contributors to colony losses (Dainat et al., 2012; Dainat and Neumann, 2013; Genersch et al., 2010; van Dooremalen et al., 2012). The microsporidium *Nosema ceranae* (*N. ceranae*) has also been implicated in regional colony losses (Botías et al., 2013; Fries et al., 2006; Higes et al., 2008; Higes et al., 2009; Martin-Hernandez et al., 2007). The threat posed by these pathogens is compounded by the fact that honey bee colonies are usually infected by several pathogens simultaneously, with *V. destructor* (together with DWV) and *Nosema* spp. being the most widespread and therefore often occurring together.

The mite *V. destructor* is an ectoparasite of honey bees (*Apis mellifera, A. cerana*) that infests honey bee colonies all over the world (for a recent review on *V. destructor* please see (Traynor et al., 2020)). The life cycle of *V. destructor* in honey bee colonies is divided into two phases, (i) the dispersal phase in which adult female mites parasitize adult bees and use the bees as a means of transport and (ii) the reproductive phase that takes place in the capped brood cell (Rosenkranz et al., 2010). For reproduction, a mature mated female mite enters a brood cell shortly before cell capping and starts laying eggs and raising her offspring soon the larva has reached the prepupal stage. For feeding, the mother mite punctures a hole in the cuticle of the developing bee. This hole is then the feeding site for the growing mite family and allows access to the pupa’s fat body, which serves as nutritional resource (Ramsey et al., 2019). Bees developing from *V. destructor* parasitized pupae show accelerated behavioral maturation, resulting in a shortened phase as nurse bees (Zanni et al., 2018), contribute less to colony productivity, and have a reduced longevity (Rosenkranz et al., 2010). Heavily mite infested colonies are characterized by an increasing rate of emerging bees which are not viable and have crippled wings. Initially, these symptoms were thought to be caused solely by mite parasitization, but it soon became clear, that *V. destructor* is an efficient virus vector (Ball, 1983; Ball, 1989) and that the crippled wings syndrome was caused by a virus, which was then named deformed wing virus (DWV) (Bailey and Ball, 1991; Bowen-Walker et al., 1999). We now know that at least four major variants of DWV exist (de Miranda et al., 2022; Martin et al., 2012; Mordecai et al., 2016; Ongus et al., 2004) and that it is the variant DWV-B that causes most of the symptoms, is more virulent than the DWV-A, and uses *V. destructor* as biological vector (Gisder et al., 2009; Gisder and Genersch, 2021; Gisder et al., 2018; McMahon et al., 2016; Posada-Florez et al., 2019; Yue and Genersch, 2005). Although *V. destructor* itself is sufficient to cause considerable damage to the parasitized pupa and the infested colony, it is the mite-vectored viruses, particularly deformed wing virus (DWV), that exacerbate the damage and link mite infestation to colony losses especially during the winter season (Dainat and Neumann, 2013; Genersch et al., 2010; Martin, 2001; Martin et al., 2012; Martin et al., 1998; McMahon et al., 2016). Microsporidia are fungal related, obligate intracellular parasites that infect many vertebrate and invertebrate host species (Keeling and Fast, 2002). Two microsporidian species infecting the adult Western honey bee *A. mellifera* are described: *Nosema apis* and *N. ceranae*. While *N. apis* is known as a honey bee-specific pathogen since more than 100 years (Zander, 1909), *N. ceranae* was originally described as pathogen of the Eastern honey bee *Apis cerana* (Fries et al., 1996), but obviously switched host several decades ago and by now is even more prevalent than *N. apis* in many *A. mellifera* populations (Botías et al., 2012; Chauzat et al., 2007; Higes et al., 2010; Klee et al., 2007; Paxton et al., 2007). However, a recent study showed that there is still no general replacement of *N. apis* by *N. ceranae*, but replacement seems to be a rather regional phenomenon (Gisder et al., 2017), presumably influenced in its dynamics by climatic conditions since *N. ceranae* spores quickly lose their infectivity when exposed to low temperatures (Fenoy et al., 2009; Gisder et al., 2010; Martin-Hernandez et al., 2009). From a clinical point of view, there is not much difference between *N. apis* and *N. ceranae*: Both pathogens frequently cause asymptomatic infections of the midgut epithelium of adult bees, but infections can also lead to diarrhea (Horchler et al., 2019). However, the factors causing the switch from asymptomatic to symptomatic infections are poorly understood (Fries, 1993; Fries, 2010). Symptomatic outbreaks of *Nosema* spp.-infections are called nosemosis and can be diagnosed by the characteristic fecal spots visible at the hive entrance and inside the hive (Horchler et al., 2019). These fecal spots contain millions of infectious spores and drive the fecal-oral transmission of the disease within the colony, as adult bees cleaning the hive of the spots ingest the spores and become infected (Bailey, 1967; Bailey and Ball, 1991). Infection in the individual adult bee host is initiated by germination of the ingested spores in the midgut lumen; germination is followed by extrusion of the polar tube, mechanical piercing of a cell by this polar tube, and injection of the sporoplasm into the cell through the polar tube (Bigliardi and Sacchi, 2001; Franzen, 2005). The reproductive cycle within the infected host cell takes about 96 hours (Gisder et al., 2011), goes through several stages (merogony, sporogony) and ends when the newly generated spores are released into the gut lumen by the bursting of the cell. These newly generated spores in the gut lumen are defecated and can infect naïve adult bees when they try to clean the hive from spore contaminated fecal spots (Bailey, 1955; Bailey, 1967; Bailey and Ball, 1991).

In 2008, the first study was published suggesting that even asymptomatic *N. ceranae* infections result in the collapse of honey bee colonies (Higes et al., 2008). Since then, numerous studies have been published demonstrating an association between *N. ceranae* infections and colony losses, but there are also many studies that failed to confirm this association (Botías et al., 2013; Fernández et al., 2012; Fries et al., 2006; Guimarães-Cestaro et al., 2020; Guzman-Novoa et al., 2011; Higes et al., 2008; Higes et al., 2009; Martin-Hernandez et al., 2007; Stevanovic et al., 2011).

Among the studies that did not observe a statistically significant association between *N. ceranae* infection and colony losses is our long-term longitudinal cohort study on honey bee health in Northeast Germany, which was initiated in 2005 and is still ongoing (Gisder et al., 2010; Gisder et al., 2017). In addition to data on colony losses during the winter season and prevalence of *Nosema* spp.-infection in spring and autumn, *V. destructor* infestation levels were also determined in each autumn over the entire study period. This unique data set enabled us to investigate the relationship between colony mortality and the two pathogens that are most commonly blamed for colony losses, *V. destructor* and *N. ceranae*. Our multivariate statistical analysis confirmed that *V. destructor* is the major cause of colony winter losses, although *N. ceranae* infections can also have deleterious effects. However, these effects are normally masked by the more severe effects of *V. destructor* on colony health and therefore only detectable in colonies that are not infested with mites or are infested at low levels. With our results we end a long-running controversy about whether or not *N. ceranae* is capable to kill entire honey bee colonies and should be considered a serious threat or even an emerging infectious disease (EID) for honey bees.

## Results

### Winter mortality

Over the last 15 years, we performed a longitudinal cohort study on honey bee colony health in the honey bee population in Northeast Germany (Genersch et al., 2010; Gisder et al., 2010; Gisder et al., 2017). We monitored between 230 and 280 colonies each year throughout the entire study period and continously collected data on winter mortality as well as on *Nosema* spp. infection status and *Varroa destructor* infestation levels in the monitored colonies. The full data set for this study comprises 3502 colonies and is a uniquely solid basis for analyzing both, the dynamics of winter colony losses and the relation between *Nosema* spp.-infection and *V. destructor*-infestation in autumn with colony losses in the following winter.

Within the study period, the winter colony loss rate varied from 4.8 % as the lowest value in the winter 2008/2009 to 26.0 % as the highest value in the winter 2016/2017. Using a linear model, we show that colony winter mortality increased by about 0.5 % per year in the studied cohort (Fig. 1A), but that this increase between 2005/2006 and 2019/2020 was not statistically significant (p-value of the F-statistic = 0.223, adjusted R^2^ = 0.043). The mean of the average winter losses was 16.31 % ± 6.56 % (mean ± SD) and therefore statistically significantly higher (p-value= 0.0023; one-sample t-test) than the empirical threshold for acceptable winter mortality of 10% (Jacques et al., 2017).

**Figure 1:**
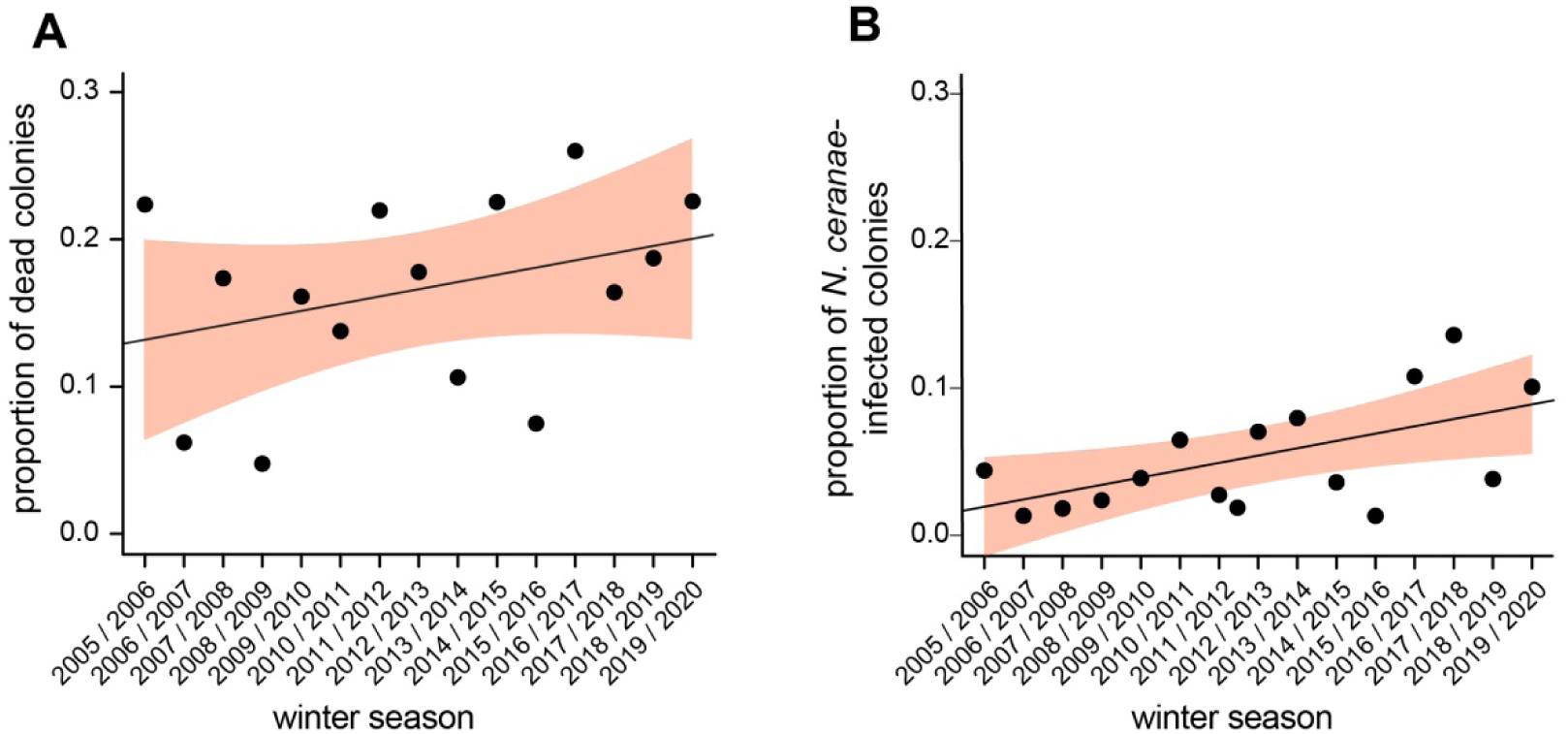
Dynamics of honey bee colony losses and *N. ceranae* infection prevalence between 2005 and 2020. Honey bee colony winter losses (A) and *N. ceranae* infection prevalence in autumn (B) were calculated over the study period from 2005 - 2020 with n=3502. Each data point represents the proportion of dead colonies in spring (A) or the prevalence of *N. ceranae*-infected colonies in autumn (B) per year. Linear regression models were calculated. Their regression lines are shown and their 95% CI (convidence interval) are highlighted in light red.

We recently demonstrated that *N. ceranae* infection prevalence in autumn showed a statistically significant increase between 2005 and 2015 (Gisder et al., 2017). We again analyzed the dynamics of *N. ceranae* infection prevalence in autumn over the entire study period of meanwhile 15 years using a linear model and demonstrate a steady increase in *N. ceranae* infection prevalence which was statistically significant (p-value of the F-statistic = 0.021, adjusted R^2^ = 0.295) and increased by about 0.5 % per year (Fig. 1B).

### Classification tree analysis

We next aimed at identifying the factors responsible for the observed increased and increasing winter losses in the monitored cohort of honey bee colonies. *V. destructor* has been identified as the main pathogen driver of winter mortality in many studies, but *N. ceranae* has also been implicated in colony losses. However, most of these studies are based on univariate analyses that observe the effect of single explanatory variables on colony mortality. Hence, data allowing to estimate the relative influence of *V. destructor* and *N. ceranae* on colony mortality when both are present in a colony, are lacking. To close this gap of knowledge, we used our data set comprising data on colony winter losses, *V. destructor* infestation levels and *Nosema* spp.-infection status and performed a classification tree analysis (Fig. 2), a tool of recursive partitioning for multivariate data exploration. This multivariate analysis aimed at identifying if *V. destructor* infestation levels or *Nosema* spp. infection status in autumn determined the fate of the colony over winter, hence, whether a colony was prone to collapse over winter or had a realistic chance to survive.

**Figure 2:**
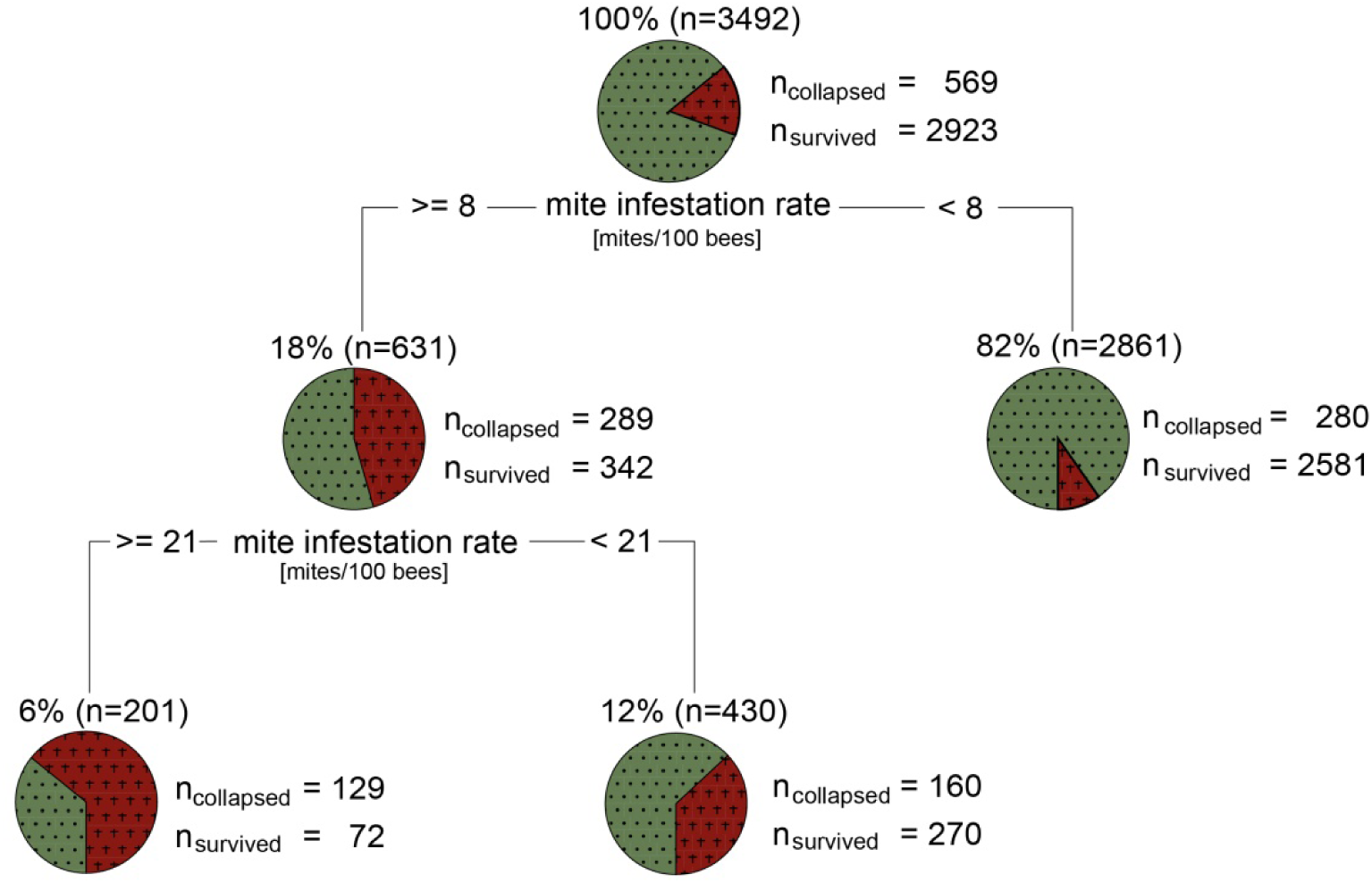
Major factors contributing to honey bee colony mortality visualized by a classification tree analysis using the R packages rpart (Therneau and Atkinson, 2019) and rattle (Williams, 2011) with standard settings.

A total of 3492 data sets was available for analysis. In the classification tree (Fig. 2), two decision points describing the fate of the colony (survival or collapse) were identified. Both were based on the mite infestation level indicating that the *Nosema* spp.-infection status of the colonies were not identified by this analysis as decisive factor for colony collapse over winter. The first decision point divided the analyzed colonies into two groups, one comprising 82% (n = 2861) of the colonies with a mite infestation rate in October of less than eight mites per 100 bees and a mortality rate of 9.8 %. The remaining 18 % of the colonies (n = 631) had an infestation level of eight or more mites per 100 bees and a mortality rate of 45.8 %. This group was further subdivided into two groups, again based on the mite infestation level. 12 % (n = 430) had an infestation level of eight or more mites but below 21 mites per 100 bees and a mortality rate of 37.2 %, whereas the other branch comprised the remaining 6 % (n = 201) of the colonies which were characterized by an infestation level of 21 or more mites per 100 bees and a mortality rate of 64.2 %. This classification tree showed convincingly that over the study period of 15 years the strongest link was between colony winter losses and *V. destructor* infestation level, but not with *N. ceranae* or *N. apis* infection.

### *N. ceranae* infection and winter mortality

The results from the classification tree analysis were in accordance with previous studies that did also not reveal any relation between colony winter losses and *Nosema* spp. infection (Genersch et al., 2010; Gisder et al., 2010; Gisder et al., 2017; Guzman-Novoa et al., 2011; Invernizzi et al., 2009; Williams et al., 2010), but contradicted other studies repeatedly showing that *N. ceranae* infections cause colony losses (Cepero et al., 2014; Higes et al., 2007; Higes et al., 2008; Higes et al., 2009; Martin-Hernandez et al., 2007). However, *Nosema* spp.-infections usually show a rather low prevalence in autumn (Bailey and Ball, 1991; Gisder et al., 2017), and hence our data set comprises only few colonies infected with *Nosema* spp. and even less infected with *N. ceranae* for each autumn. We, therefore, speculated that the number of *Nosema* spp. infected colonies per year might be too low to see a relation between winter mortality and *Nosema* spp. infection and decided to generate higher numbers for *Nosema* infected colonies by summing the numbers for the individual years starting with autumn/winter 2005/2006 and ending with the sum of autumn/winter 2005/2006 up to autumn/winter 2019/2020 (Table 1). Forming these cumulative data subsets and examining time periods instead of annual values, resulted in larger group sizes (n’s) and overall larger numbers of *Nosema* spp. infections, which made statistical analyses more robust (Table 1). We used a Chi-squared test to calculate the statistical relationship between infection status and winter mortality and indeed, the analysis started to become significant (p-value < 0.05) when more than 11 years (2005/2006 up to 2016/2017 resulting in more than 221 *Nosema* spp.-infected colonies among the analyzed 3492 colonies) were considered (Table 1, Fig. 3). This was the first time that we were able to show a statistically significant impact of *Nosema* infection on colony losses.

**Table 1:** Cumulative data subsets for determining the effects of *Nosema* spp.-infection in the autumn on honeybee colony losses in the following winter.

| Cumulative overwintering period | Total number of colonies | Surviving colonies |  | Collapsed colonies |  | p<sup>a</sup> |
| --- | --- | --- | --- | --- | --- | --- |
|  |  | *Nosema* positive | *Nosema* negative | *Nosema* positive | *Nosema* negative |  |
| 2005 | 237 | 21 | 163 | 9 | 44 | 0.28 |
| 2005 - 2006 | 463 | 40 | 356 | 10 | 57 | 0.24 |
| 2005 - 2007 | 682 | 51 | 526 | 14 | 91 | 0.15 |
| 2005 - 2008 | 892 | 62 | 715 | 14 | 101 | 0.13 |
| 2005 - 2009 | 1072 | 75 | 853 | 17 | 127 | 0.14 |
| 2005 - 2010 | 1319 | 96 | 1045 | 22 | 156 | 0.09 |
| 2005 - 2011 | 1574 | 111 | 1229 | 26 | 208 | 0.16 |
| 2005 - 2012 | 1844 | 133 | 1429 | 29 | 253 | 0.33 |
| 2005 - 2013 | 2070 | 166 | 1598 | 35 | 271 | 0.27 |
| 2005 - 2014 | 2292 | 175 | 1761 | 38 | 318 | 0.33 |
| 2005 - 2015 | 2519 | 182 | 1964 | 39 | 334 | 0.21 |
| 2005 - 2016 | 2769 | 203 | 2128 | 53 | 385 | 0.02 |
| 2005 - 2017 | 3019 | 234 | 2306 | 66 | 413 | < 0.01 |
| 2005 - 2018 | 3254 | 240 | 2491 | 73 | 450 | < 0.001 |
| 2005 - 2019 | 3502 | 267 | 2656 | 74 | 505 | <0.01 |

<sup>a</sup> Determined by the $\chi^2$ test

**Figure 3:**
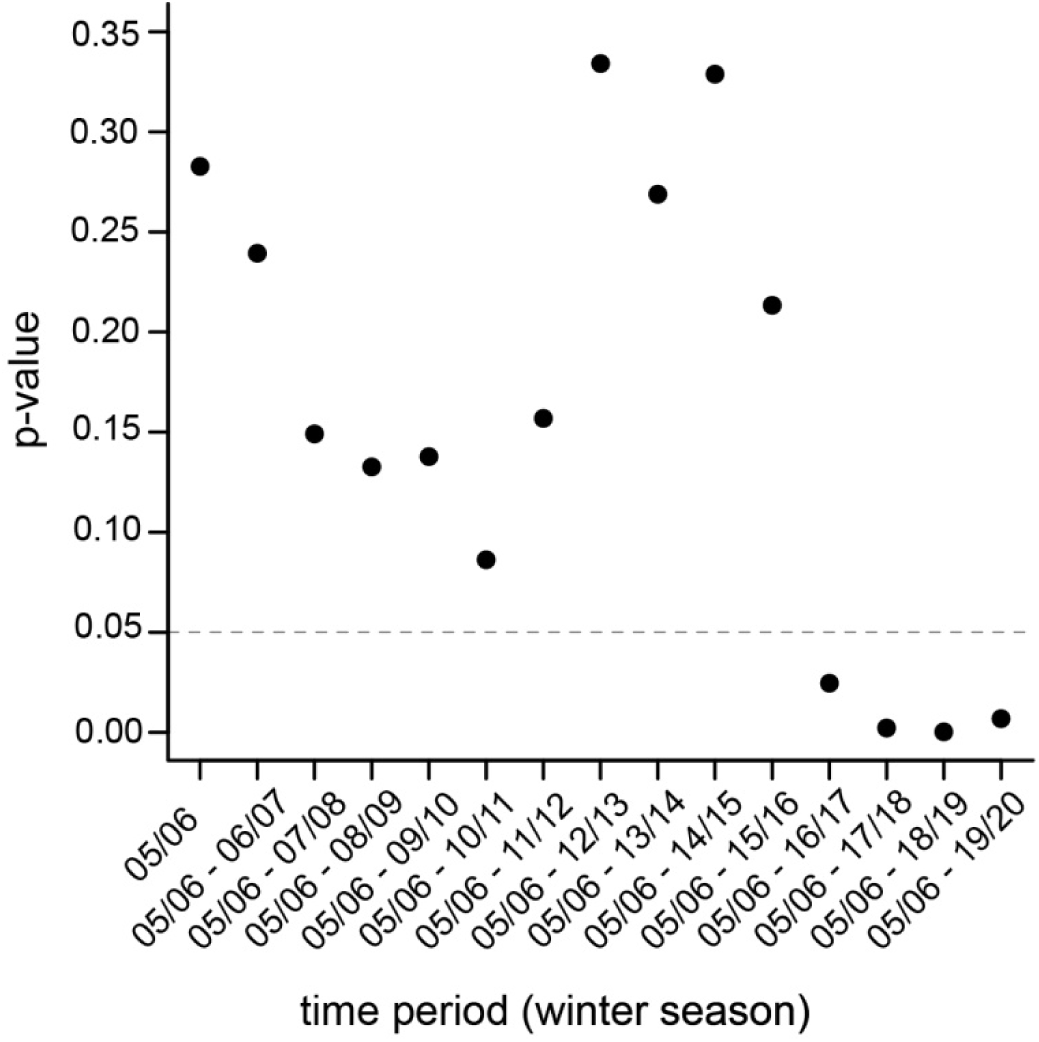
Calculated p-values for the relation between colony losses and Nosema-infection status using cumulated data for the numbers of infected clonies. Each data point represents the cumulative number of infected colonies summed over the time window indicated on the x-axis, while the y-axis gives the respective p-value of the Chi-squared test.

To further support and analyze this impact, we performed a Chi-squared test giving Pearson residuals using the data cumulated over the entire study period (2005/2006 to 2019/2020) for *Nosema* spp. infection (Fig. 4A) and for *N. apis, N. ceranae* and mixed infections separately (Fig. 4B). The black lines in the associations plots (Fig. 4) represent the expected values for the categories “alive” or “dead” in relation to the infection categories negative or positive (Fig. 4A) or negative and positive for *N. apis*-, *N. ceranae*-or co-infections (Fig. 4B). Overrepresented categories are represented by rectangles above the base line, while the rectangles for underrepresented categories are below the base line. Categories with Pearson residuals above 2.0 are shown in blue and appear only for the combinations “dead” and “positive for *Nosema* spp.-infection” (Fig. 4A) or “dead” and “positive for *N. ceranae* infection” (Fig. 4B). These results confirmed that the relation between Nosema infection and winter colony losses was statistically significant (p-value = 0.007). However, this was only true for *N. ceranae*-infections (p-value = 0.002); *N. apis*-or co-infections did not significantly contribute to winter losses suggesting that *N. ceranae* is indeed more virulent at colony level than *N. apis*.

**Figure 4:**
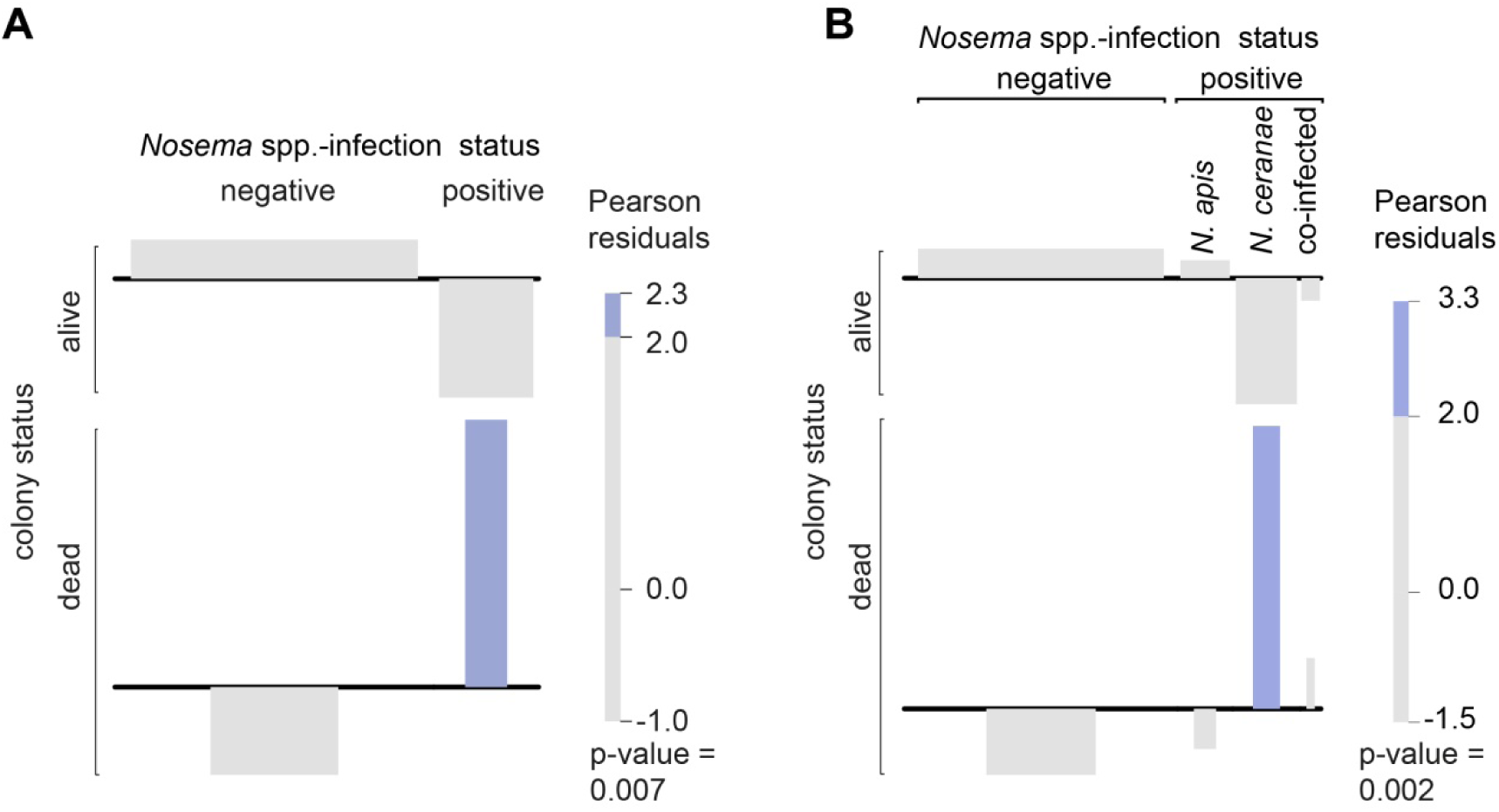
Contingency table analysis using association plots based on Chi-squared testing of the association between *Nosema* spp.-infection and colony mortality (A) and *N. apis*-, *N. ceranae* and co-infections and colony mortality (B). Overrepresented entries (dead/positive in A or dead/*N. ceranae* in B) are shown in blue.

### Statistical significance vs. biological relevance

P-values are a measure of statistical significance, but are insufficient to show biological relevance. To analyze the biological relevance of our results, we therefore used the effect size measure Cohen’s ω, which is applicable for two times two and larger contingency tables and is considered a measure of relevance with values above 0.1, 0.3, and 0.5 indicating a small, medium, and large effect size, respectively (Cohen, 1988). Calculating Cohen’s ω for our data set and the relation between colony losses and *Nosema* spp.-or *N. ceranae*-infection revealed an effect size below 0.1, hence, a less than small effect (Fig. 5A). These results indicated that although we showed a statistically significant relationship between colony losses and *N. ceranae*-infection, these relationships are of minor or no biological relevance supporting the results of the classification tree analysis (Fig. 2) identifying *V. destructor* as main factor in colony losses.

**Figure 5:**
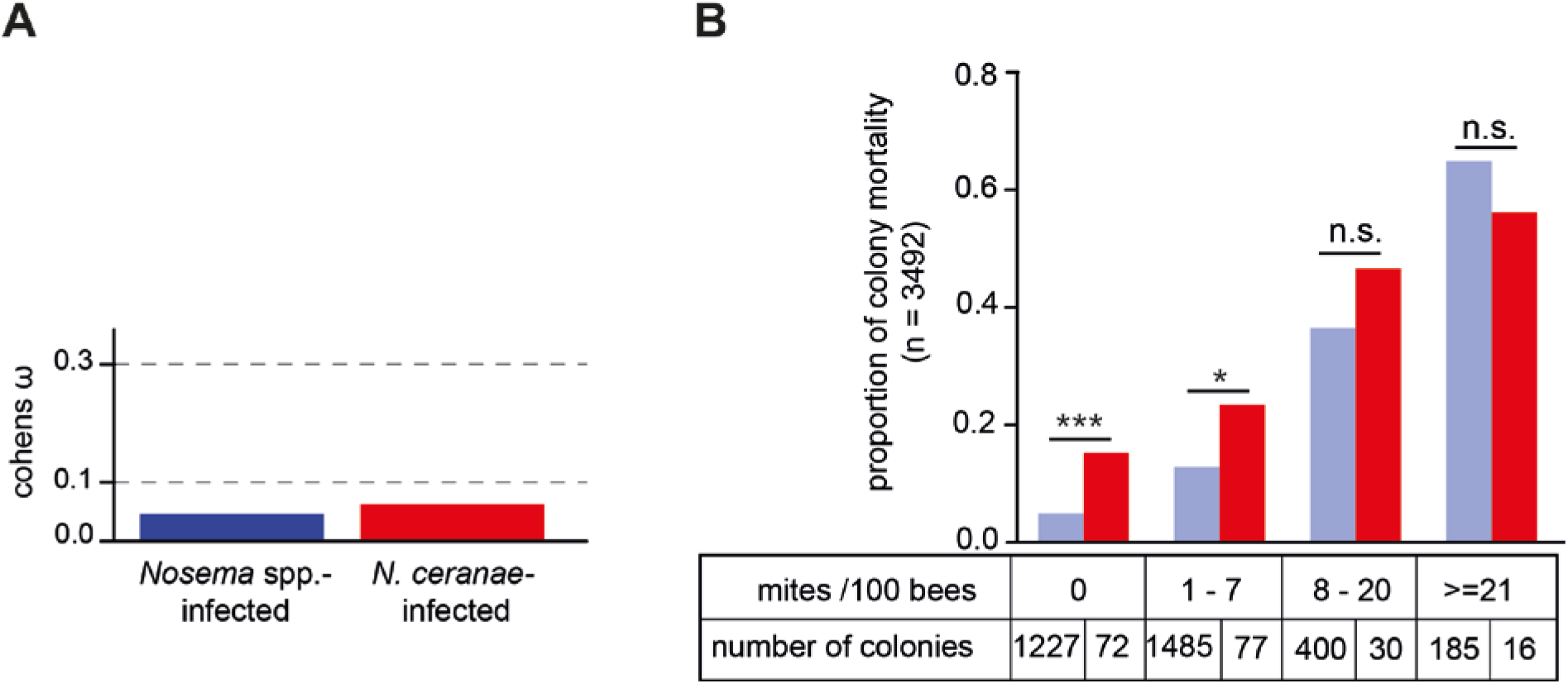
Biological relevance of *N. ceranae* infections. (A) Bar plot showing the effect size Cohen’s ω for *Nosema* spp.- and *N. ceranae*-infected colonies on colony mortality. Dashed lines indicate the conventional definition of Cohen’s ω with a value for ω between 0.1 and 0.3 as small, between 0.3 and 0.5 as medium and above 0.5 as large effect (Cohen, 1988). (B) Bar plot showing the relation between colony mortality, mite infestation level categories and *N. ceranae*-infection status. Significance levels are indicated by asterisks (n.s., p ≥ 0.05; significantly different: *, 0.05 < p < 0.01; **, 0.01 < p < 0.001; ***, 0.001 > p).

We therefore reanalyzed the data, but this time we considered the *V. destructor* infestation rate of the colonies by analyzing the mortality rate of colonies infected and not infected with *N. ceranae* within the mite infestation categories as defined by the classification tree (Fig. 2). This analysis revealed that *N. ceranae* infection only contributed siginficantly to colony mortality when the colonies did not harbor detectable *V. destructor* mites or very few mites (1-7 per 100 bees) in October. Only in these 149 out of 3474 colonies, *N. ceranae* was significantly correlated with winter mortality (Fig. 5B).

## Discussion

Honey bee colony losses and the quest for abiotic and biotic factors causing them are hot topics in the field of bee research since two decades. While it is widely accepted and unequivocally substantiated by many studies, that the ectoparasitic mite *V. destructor* and the viruses vectored by the mite play a key role in winter colony losses, the role of *N. ceranae* infections is less clear. There are studies clearly showing detrimental effects of *N. ceranae* infection on honey bee colonies (Botías et al., 2013; Fries et al., 2006; Higes et al., 2008; Higes et al., 2009; Martin-Hernandez et al., 2007). But there are also numerous monitoring studies that fail to observe such an effect (Fernández et al., 2012; Gisder et al., 2010; Gisder et al., 2017; Guimarães-Cestaro et al., 2020; Stevanovic et al., 2011). One possible reason for this could be that in monitoring studies the damage caused by the almost ubiquitous infestation of colonies with *V. destructor* masks the effects caused by pathogens with rather low prevalance such as *N. ceranae*. This masking effect is difficult to see through because most observational studies on colony losses include too few colonies and are conducted over too short a time period to observe statistically significant associations for low-prevalence pathogens. Such monitoring studies usually collect data for multiple pathogens enabling to look for the interaction between co-existing pathogens. For example, *N. ceranae*-infections in spring have been shown to correlate statistically significantly with an increased prevalence of *Ascosphaera apis* infections and higher levels of *V. destructor* infestation in summer (Hedtke et al., 2011). However, there are only few data on the relative impact of individual pathogens on colony mortality, although it is widely accepted that colony collapse is a multifactorial process, often likely involving multiple pathogens.

Our data on pathogen load and winter colony mortality, collected continuously over 15 years from a relatively stable cohort of about 25 apiaries contributing ten colonies each is a unique resource to study the role of *N. ceranae* on colony mortality, especially the relative impact of *N. ceranae* infections on overwintering success of honey bee colonies, most of which were concurrently infested by *V. destructor*. With more than 3000 data sets collected over 15 years, we were able to confirm that big data sets and long study durations are key for robust analyses: Only by summing the data for more than 11 years did a statistically significant association between *N. ceranae-*, but not *N. apis*-infection in the autumn and colony losses the following winter become evident. This result indicated that at colony level *N. ceranae* is indeed more virulent than *N. apis* confirming previous studies on *N. ceranae*-induced colony losses (Botías et al., 2013; Fries et al., 2006; Higes et al., 2008; Higes et al., 2009; Martin-Hernandez et al., 2007). No such relation between *N. ceranae*-infections and colony losses was observed when the data were analyzed year by year (Gisder et al., 2010; Gisder et al., 2017) because the prevalence of *Nosema* spp.-infections in autumn and the mortality rate among these colonies are usually too low for statistical significance explaining previous studies that did not support *N. ceranae*-induced colony losses (Fernández et al., 2012; Gisder et al., 2010; Gisder et al., 2017; Guimarães-Cestaro et al., 2020; Stevanovic et al., 2011).

A statistically significant association between *N. ceranae* infection and colony losses was in accordance with studies that had suggested an increased virulence of *N. ceranae* compared to *N. apis* and the abilty of *N. ceranae* to cause the collapse of entire colonies (Botías et al., 2013; Fries et al., 2006; Higes et al., 2008; Higes et al., 2009; Martin-Hernandez et al., 2007). However, our multivariate data exploration via classification tree analysis had identified *V. destructor* infestation as the main variable explaining colony losses in our cohort. A result that is also in accordance with many other studies clearly linking mite infestation to winter colony mortality (Dainat and Neumann, 2013; Genersch et al., 2010; Martin, 2001; Martin et al., 2012; Martin et al., 1998; McMahon et al., 2016). Only with a mite infestation rate in October of less than eight mites per 100 bees, an acceptable winter mortality rate below 10 % (Jacques et al., 2017) can be reached. In the monitored cohort this was the case for 82 % of the colonies over the entire duration of the study, leaving 18 % of the cohort contributing to inacceptably elevated colony losses. Since classification tree analysis is the method of choice for determining biologically relevant factors and the classification tree identified mite infestation as the only relevant factor, it was not surprising that determination of Cohen’s ω confirmed that the biological relevance of *N. ceranae* infection for colony losses is low despite the statistical significance of this association. This clearly shows that for biological questions the focus should rather not soley lay on statistical significance but more on the effect size and biological relevance.

We showed that *N. ceranae* infection contributed to colony losses only in those colonies that were not or low infested by *V. destructor* in autumn. As long as *V. destructor* infestation is the dominating health problem in honey bee colonies and *N. ceranae* prevalence is low, *N. ceranae* can be classified as pathogen causing little concern, because its role in colony losses is marginal. However, the situation might change when the prevalence of *N. ceranae* reaches a critical point. Since the increase in *N. ceranae* prevalence is continuing ((Gisder et al., 2017) and this study), it is only a question of time when this point will be reached. Monitoring not only mite infestation levels in colonies but also *N. ceranae* infection prevalence in honey bee populations is therefore advisable. Hence, we will continue our study and continuously calculate the effect size of *N. ceranae* infection on colony losses to determine the critical prevalence of *N. ceranae* in a honey bee population.

Remarkable is that in an early report on colony collapse due to *N. ceranae* (Higes et al., 2009) it was explicitly pointed out that mites were absent in all samples indicating a very low number or even the total absence of *V. destructor* in these collapsed colonies due to efficient mite control. The absence of *V. destructor* and concomitant presence of *N. ceranae* was the most convincing argument for *N. ceranae* being the cause of colony collapse in the reported case. Moreover, for experimentally demonstrating *N. ceranae*-induced colony collapse, the colonies needed to be tightly controlled for *V. destructor* infestation (Higes et al., 2008). These studies corroborate our results which indicate that *N. ceranae*-induced colony collapse only becomes evident in (nearly) mite-free colonies. Hence, the more efficient the mite control is, the more likely it is that *N. ceranae* induced damages become detectable. But again, as long as *N. ceranae* prevalence is low this does not pose a serious threat because *N. ceranae* is still not a highly virulent pathogen.

## Material and Methods

### Bee Samples, Field Survey

The data set of this study comprises samples which were collected from autumn 2005 to spring 2020 in the course of a 15 year longitudinal cohort-study on *Nosema* spp. epidemiology and honey bee health (Genersch et al., 2010; Gisder et al., 2010; Gisder et al., 2017). Honey bee samples were collected in autumn and colonies were checked for their survival in spring of the respective overwintering period (weeks 36 to week 14 of the following year) from about 23 apiaries which were located in Northeast-Germany (Fig. 6). Briefly, apiaries participated with ten so called “monitoring colonies” each. Apiaries or monitoring colonies that dropped out during the study period were substituted by adequate replacement. Hence, more than half of the apiaries (14 of ∼23) participated for more than 9 years and 5 of them even for the entire duration of the study, *i*.*e*. 15 years. From at least 19 apiaries, samples were provided over a time period of consecutive 5-11 years (Fig. 6). This resulted in an annual mean of 23.4 ± 2.26 (mean ± SD) apiaries with 9.77 ± 1.25 (mean ± SD) colonies each, giving an overall count of n = 3502 sampled monitoring colonies which provide the basis of our analyses.

**Figure 6:**
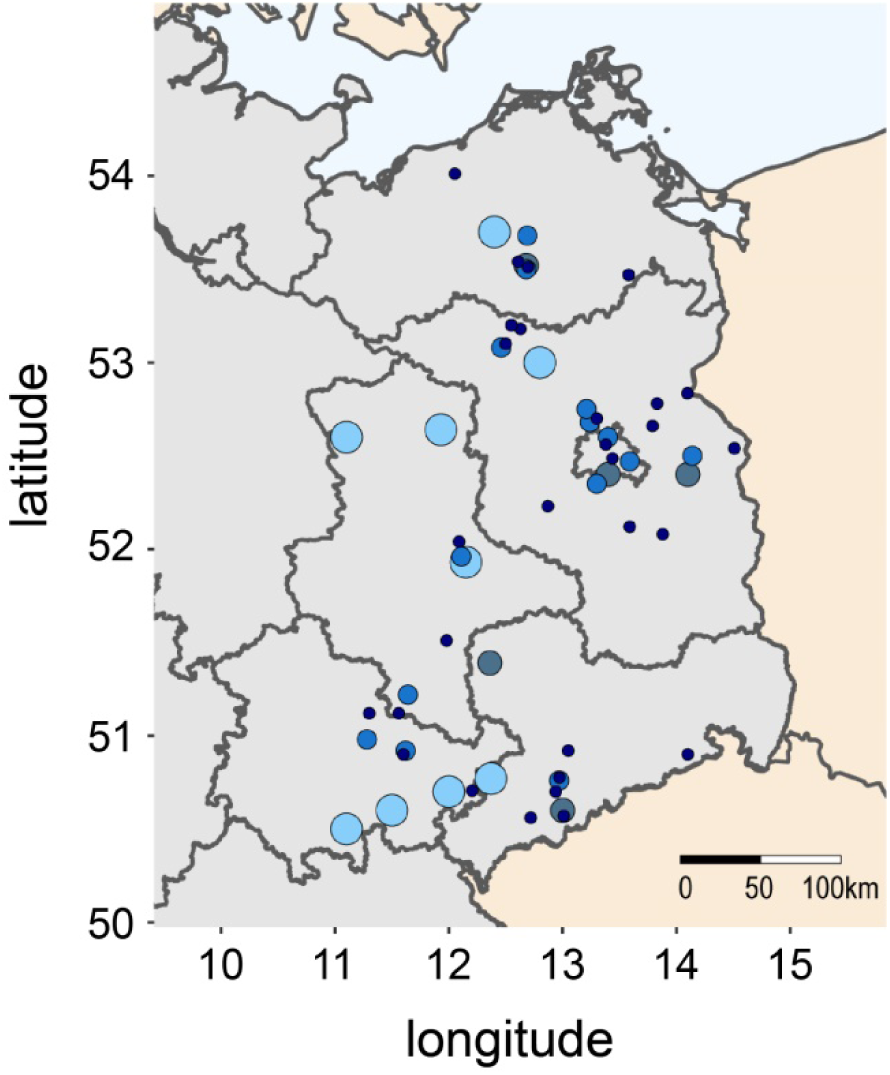
Map section of Northeast Germany showing the location of the apiaries which participated in the study. The size and color of the circles represent the number of years for which data are available for each apiary (light blue, 12 - 15 years; grey-blue, 9 - 11 years; royal blue, 5 - 8 years; dark blue, 1 - 4 years)

Sampling of bees was performed essentially as already described (Gisder et al., 2010; Gisder et al., 2017). Briefly, between calendar week 36 and 38 (late September/ beginning of October), about 300 in-hive honey bees were sampled from a super above the queen excluder (Fries et al., 2013) from each monitoring colony. Bee samples were stored at -20 °C until further analysis.

### Determination of mite infestation levels

For determining the mite infestation level, *V. destructor* mites were washed from about 150 sampled bees following a standard protocol using a detergent solution (Dietemann et al., 2013). Briefly, the frozen bees were covered with soap and water in a jar. Subsequently, the jar was shaken for 20 seconds and emptied into two stacked sieves with a white nylon cloth between them. All mites were rinsed with plenty of water under high pressure through the upper sieve which has larger aperture (3-4 mm) not allowing bees to pass. All mites were collected on the cloth in the second sieve, which has smaller aperture (<0.5 mm) that no mite fits through. To gain the mite infestation level in %, the number of counted mites was divided by the number of sampled and washed bees multiplied by 100. Unfortunately, mite infestation rate could not be determined in one apiary (ten colonies) in the first year (2005 / 2006) because of an insufficient number of honey bees available. Therefore, the dataset for mite infestation level had to be reduced from 3502 to 3492.

### Diagnosis of *Nosema* spp. infection and molecular species differentiation

Diagnosis of *Nosema* spp. infections was performed as already described (Genersch et al., 2010; Gisder et al., 2010; Gisder et al., 2017) and in accordance with the “Manual of Standards for Diagnostics and Vaccines” published by the Office International des Epizooties (OIE), the World Organization for Animal Health (Anonymous, 2021). In short, per colony 20 pooled bee abdomens were homogenized and microscopically examined for the presence of spores. Infection levels were determined by counting the number of *Nosema* spp. spores per view field (three technical replicates each) in a hemocytometer (Neubauer-improved, VWR, Darmstadt, Germany) using an inverse microscope (VWR, Darmstadt, Germany) with 100 × magnification. For classification of the infection levels, standard categories were used (Anonymous, 2021): 0 (no spores), 1 (1-10 spores), 2 (11-100 spores), and 3 (more than 100 spores).

*Nosema* spp.-positive samples were subjected to further molecular species differentiation either via PCR-amplification of a conserved region of the16S rRNA gene followed by RFLP (restriction fragment length polymorphism) analysis of this amplicon (Gisder et al., 2010; Klee et al., 2007) or via a species-specific duplex PCR-protocol taking advantage of species-specific sequence differences in the highly conserved gene coding for the DNA-dependent RNA polymerase II largest subunit (Gisder and Genersch, 2013). Molecular differentiation enabled the distinction between single infections with either *N. apis* or *N. ceranae*, or co-infections where both infections are present at the same time (Table 2).

**Table 2:**
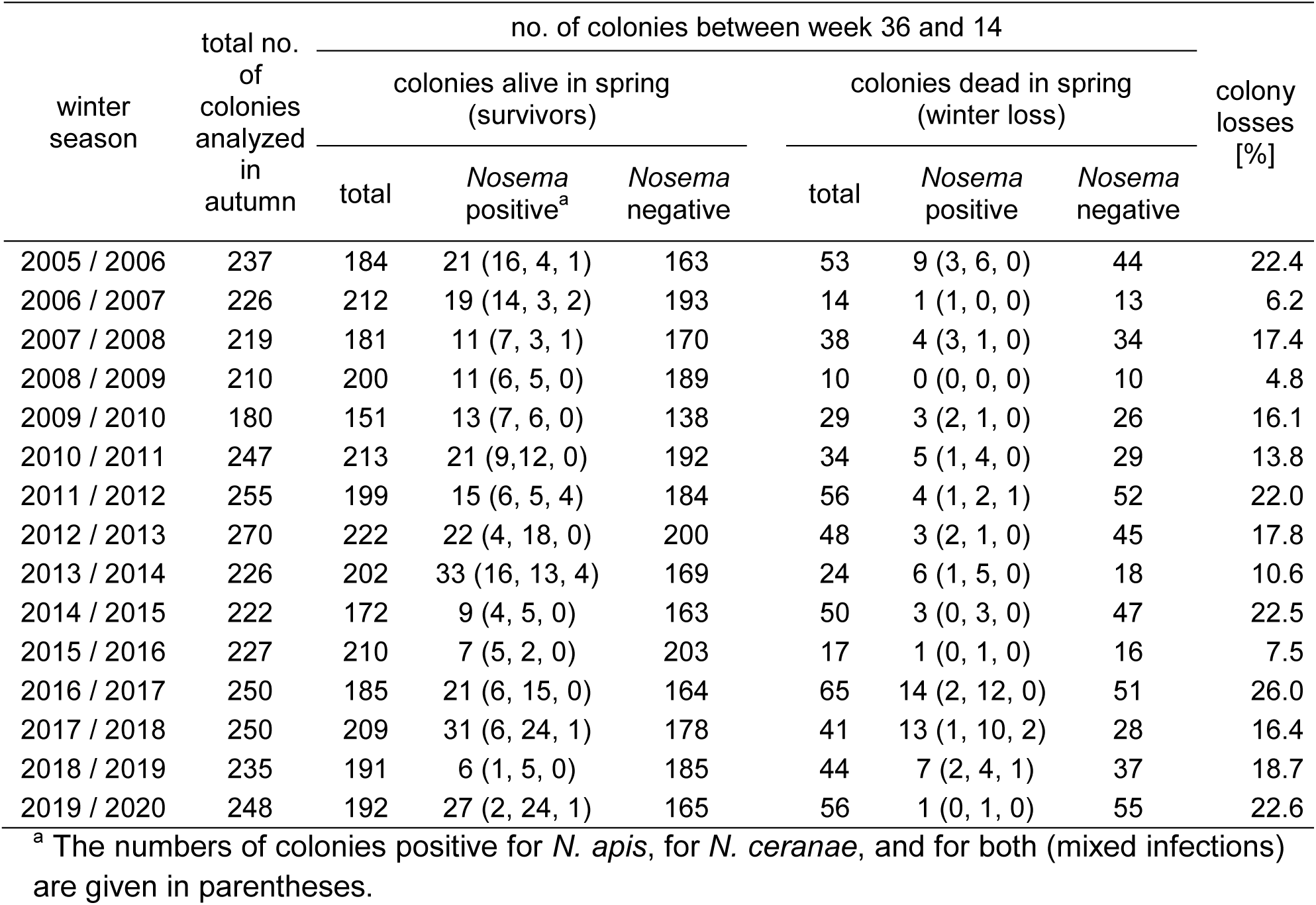
Data on *Nosema* spp. epidemiology (prevalence of *Nosema* spp.-, *N. apis*-, *N. ceranae*- and co-infections) and winter losses collected between autumn 2005 and spring 2020

| winter season | total no. of colonies analyzed in autumn | no. of colonies between week 36 and 14 |  |  |  |  |  | colony losses [%] |
| --- | --- | --- | --- | --- | --- | --- | --- | --- |
|  |  | colonies alive in spring (survivors) |  |  | colonies dead in spring (winter loss) |  |  |  |
|  |  | total | *Nosema* positive<sup>a</sup> | *Nosema* negative | total | *Nosema* positive | *Nosema* negative |  |
| 2005 / 2006 | 237 | 184 | 21 (16, 4, 1) | 163 | 53 | 9 (3, 6, 0) | 44 | 22.4 |
| 2006 / 2007 | 226 | 212 | 19 (14, 3, 2) | 193 | 14 | 1 (1, 0, 0) | 13 | 6.2 |
| 2007 / 2008 | 219 | 181 | 11 (7, 3, 1) | 170 | 38 | 4 (3, 1, 0) | 34 | 17.4 |
| 2008 / 2009 | 210 | 200 | 11 (6, 5, 0) | 189 | 10 | 0 (0, 0, 0) | 10 | 4.8 |
| 2009 / 2010 | 180 | 151 | 13 (7, 6, 0) | 138 | 29 | 3 (2, 1, 0) | 26 | 16.1 |
| 2010 / 2011 | 247 | 213 | 21 (9,12, 0) | 192 | 34 | 5 (1, 4, 0) | 29 | 13.8 |
| 2011 / 2012 | 255 | 199 | 15 (6, 5, 4) | 184 | 56 | 4 (1, 2, 1) | 52 | 22.0 |
| 2012 / 2013 | 270 | 222 | 22 (4, 18, 0) | 200 | 48 | 3 (2, 1, 0) | 45 | 17.8 |
| 2013 / 2014 | 226 | 202 | 33 (16, 13, 4) | 169 | 24 | 6 (1, 5, 0) | 18 | 10.6 |
| 2014 / 2015 | 222 | 172 | 9 (4, 5, 0) | 163 | 50 | 3 (0, 3, 0) | 47 | 22.5 |
| 2015 / 2016 | 227 | 210 | 7 (5, 2, 0) | 203 | 17 | 1 (0, 1, 0) | 16 | 7.5 |
| 2016 / 2017 | 250 | 185 | 21 (6, 15, 0) | 164 | 65 | 14 (2, 12, 0) | 51 | 26.0 |
| 2017 / 2018 | 250 | 209 | 31 (6, 24, 1) | 178 | 41 | 13 (1, 10, 2) | 28 | 16.4 |
| 2018 / 2019 | 235 | 191 | 6 (1, 5, 0) | 185 | 44 | 7 (2, 4, 1) | 37 | 18.7 |
| 2019 / 2020 | 248 | 192 | 27 (2, 24, 1) | 165 | 56 | 1 (0, 1, 0) | 55 | 22.6 |

<sup>a</sup> The numbers of colonies positive for *N. apis*, for *N. ceranae*, and for both (mixed infections) are given in parentheses.

## Statistical Analysis

Data was curated, transformed and presented in spreadsheets for the analysis with the statistic software R (R Development Core Team, 2021) using the package *openxlsx* (Schauberger et al., 2021). Data for the map section of Northeast Germany (Fig. 6) were obtained from the website http://www.gadm.org (Hijmans et al., 2016). The map was created with R by the use of the following packages: *rnaturalearth* (South, 2017), *raster* (Hijmans, 2022), *ggplot2* (Wickham, 2016), *sf* (Pebesma, 2018), *sp* (Bivand et al., 2013; Pebesma and Bivand, 2005), *rgeos* (Bivand and Rundel, 2021), and *reshape* (Wickham, 2007).

For the winter losses we performed a linear regression model (using R base package *stats*) and calculated its regression line, its slope, the adjusted R^2^ and the F-statistic. To describe and visualize which variable(s) has (have) the largest share in honey bee colony mortality, we took advantage of classification tree analysis (decision trees), a tool of recursive partitioning for multivariate data exploration. We constructed a tree with default settings using *rpart* (Therneau and Atkinson, 2019) and *rattle* (Williams, 2011).

To understand the role of *N. ceranae* infection in more detail, we looked at the *Nosema* spp. infection frequencies using contingency tables and performed chi-squared calculation. The results were plotted in two-way association plots created by *vcd* (Meyer et al., 2006; Meyer et al., 2007; Meyer et al., 2020). The obtained results of the significance statistic were further assessed by strength statistic using the effect size index Cohen’s ω (Cohen, 1988), which is an index for the biological effect size between two categorical variables. Cohen’s ω is calculated as follows:

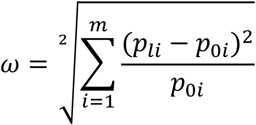

p_li_ = the proportion in cell i posited by the alternate hypothesis and reflects the effect for that cell; p_0i_ = the proportion in cell i posited by the null hypothesis; m = number of cells [(Cohen, 1988) p. 216 formula 7.2.1]. For the interpretation of Cohen’s ω in terms of effect size, there is a framework of conventional definition saying that 0.1 ≤ ω <0.3 is a small, 0.3 ≤ ω <0.5 is a medium and ω ≥ 0.5 is a large effect size [(Cohen, 1988) chapter 7.3 p.227].

## Acknowledgements

This study was supported by the Ministries responsible for Agriculture of the German Federal States of Brandenburg, Saxonia-Anhalt, Saxonia, Thuringia, and of the Senate of Berlin, Germany, by a grant from the German Ministry of Agriculture through the Bundesanstalt für Landwirtschaft und Ernährung (BLE, grant no. 2816SE004), and by a grant of the German Research Foundation (DFG, GRK2046).

